# Theoretical framework and experimental solution for the air-water interface adsorption problem in cryoEM

**DOI:** 10.1101/2023.05.23.541984

**Authors:** Joon S. Kang, Xueting Zhou, Yun-Tao Liu, Kaituo Wang, Z. Hong Zhou

## Abstract

As cryogenic electron microscopy (cryoEM) gains traction in the structural biology community as a method of choice for determining atomic structures of biological complexes, it has been increasingly recognized that many complexes that behave well under conventional negative-stain electron microscopy tend to have preferential orientation, aggregate or simply mysteriously “disappear” on cryoEM grids, but the reasons for such misbehavior are not well understood, limiting systematic approaches to solving the problem. Here, we have developed a theoretical formulation that explains these observations. Our formulation predicts that all particles migrate to the air-water interface (AWI) to lower the total potential surface energy — rationalizing the use of surfactant, which is a direct solution to reducing the surface tension of the aqueous solution. By conducting cryogenic electron tomography (cryoET) with the widely-tested sample, GroEL, we demonstrate that, in a standard buffer solution, nearly all particles migrate to the AWI. Gradual reduction of the surface tension by introducing surfactants decreased the percentage of particles exposed to the surface. By conducting single-particle cryoEM, we confirm that applicable surfactants do not damage the biological complex, thus suggesting that they might offer a practical, simple, and general solution to the problem for high-resolution cryoEM. Application of this solution to a real-world AWI adsorption problem with a more challenging membrane protein, namely, the ClC-1 channel, has led to its first near-atomic structure using cryoEM.

## 1. Introduction

Cryogenic electron microscopy (cryoEM) has become a tool of choice for determining the atomic structures of biological complexes. The general strategy in sample preparation involves 3 steps: First, the sample is placed on continuous carbon film, negatively stained, and evaluated by transmission electron microscopy (TEM) to optimize the purity and concentration of the sample. Second, a droplet of the optimized sample is applied to various cryoEM grids (*e*.*g*., holey carbon or gold foil grids, graphene film grids, lacey carbon film grids, *etc*.) and blotted with filter paper to create a thin layer of liquid containing the sample. Third, the blotted grid is plunged into liquid ethane to flash-freeze the thin layer of liquid, embedding the sample in vitreous ice (Dubochet et al., 1988). After these steps, one often observes, surprisingly and disappointedly, aggregation and deformation of the embedded particles of the sample (*e*.*g*., Fig. 1) at the air-water interface (AWI) (Taylor and Glaeser, 2008). Cryogenic electron tomography (cryoET) revealed that a broad range of macromolecular complexes was adsorbed at the AWI in cryoEM grid holes when these standard grid preparation steps were used (Noble et al., 2018b). This problem, also known as the AWI adsorption phenomenon, causes uneven distribution and preferred orientations of the particles, resulting in directional resolution anisotropy (Lyumkis, 2019) — a major obstacle in high-resolution cryoEM structure determination (Bai et al., 2013; D’Imprima et al., 2019). This AWI adsorption phenomenon was reported decades ago (Graham and Phillips, 1979; MacRitchie, 1985; Narsimhan and Uraizee, 1992), affecting membrane and non-membrane proteins (Noble et al., 2018b). It has been proposed that this phenomenon results from the particles’ diffusion to the AWI due to their exposed hydrophobic regions (Glaeser and Han, 2017). However, a complete understanding and elimination of this phenomenon remain elusive.

**Figure 1.**
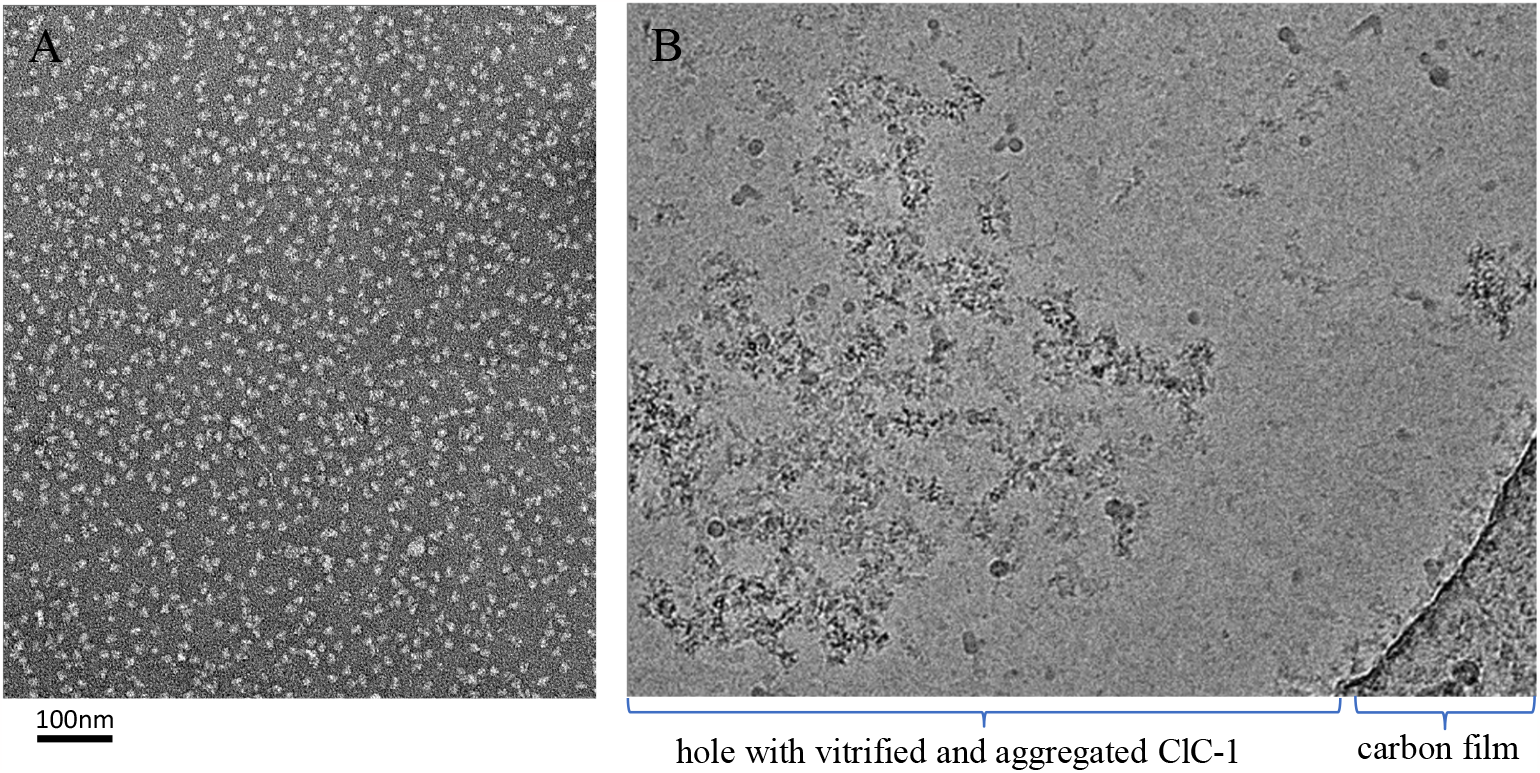
Example of the AWI adsorption problem in cryoEM. (A) Negative stain EM micrograph of purified human ClC-1 protein particles, showing intact particles and optimal particle distribution prior to vitrification. (B) CryoEM micrograph of frozen-hydrated human ClC-1 protein particles across holey carbon film, showing the typical aggregation and deformation problem occurring at the AWI in the absence of adequate surfactant.

Despite the lack of understanding, numerous attempts have been made to develop practical methods to address the problem. For example, using a faster plunge device (such as Spotiton), whose elapsed time between applying the sample to grids and plunging into liquid ethane is in the order of 100 ms (much less than the traditional plunge of more than 1s), can relieve preferred orientations problem caused by the AWI phenomenon (Jain et al., 2012; Noble et al., 2018a).Using continuous thin layers of amorphous carbon (Williams and Glaeser, 1972) or graphene (D’Imprima et al., 2019; Pantelic et al., 2011; Russo and Passmore, 2014) also alleviated the AWI adsorption problem. Although these methods may improve the cryoEM result, they still have limitations. For one, using a fast nano-dispenser does not provide direct evidence of eliminating AWI absorption problems (Noble et al., 2018a). Using a thin layer of amorphous carbon as a support layer on cryoEM grids can adsorb and thus immobilize some particles, but it also adds background noise to cryoEM images (Russo and Passmore, 2016). A more widely used method is to include surfactants or detergents in the sample solution to improve particle preservation on cryoEM grids, especially for membrane proteins. For example, zwitter-ionic detergent (CHAPSO) has been reported to reduce the AWI adsorption of bacterial RNA polymerase (Chen et al., 2019) and human erythrocyte catalase (Chen et al, 2022) and alleviate the preferred orientation problem. Unfortunately, a particular surfactant may work well for one sample type but not for others. Therefore, a deeper understanding of the mechanism of AWI is needed to better use surfactants in cryoEM sample preparation.

In this study, we report a theoretical framework to explain the AWI adsorption phenomenon in the context of surface energy and experimentally examined the effects of different surfactants on cryoEM sample preparation with the standard test sample, GroEL, using cryoET and cryoEM. According to this framework, particles migrate to AWI to decrease the overall area of the liquid exposed to the air, thereby minimizing the overall potential energy of the system. This framework rationalizes using surfactants to alleviate the AWI problem: They lower the surface energy of an aqueous solution, decreasing the overall potential energy and the particles’ tendency to migrate to AWI. To experimentally test the theory’s predictability, we used cryoET to visualize particle distribution across the sample depth on the cryoEM grid from one AWI to the other. Our cryoET results demonstrate the effectiveness of three surfactants [NP40, DDM, and fluorinated fos-choline 8 (FFC8)] against the AWI adsorption problem. We also conducted single-particle cryoEM and 3D reconstruction with GroEL to demonstrate that these surfactants did not adversely affect the protein structures at near-atomic resolution. We further show the effectiveness of FFC8 in drastically improving particle distribution and generating a near-atomic resolution structure of a more challenging membrane protein, the ClC-1 channel.

## 2. Results

### 2.1 Theoretical formulation of the air-water interface problem in cryoEM

All matter, except for ideal gases, is held together by molecules that have varying degrees of attraction to one another. In the bulk of matter, the intermolecular forces equilibrate each other. However, at the surface of matter, molecules are not fully surrounded by their neighbors, which results in a net inward force pointing into the bulk (Hauner et al., 2017). This net inward force increases as the exposed surface area of matter increases. To achieve equilibrium, work must be done to counterbalance this net inward force; The greater the exposed surface area of matter, the greater the work to counterbalance the net inward force. The ratio between the work and surface area, or work per unit surface area, can be represented by (γ):

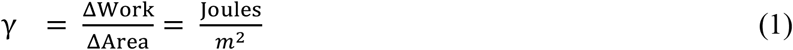

which is also known simply as *surface energy* in the field of surface science and fluid dynamics. Surface energy is due to the attraction of electrical charges around the molecules. Surface energy is then defined as the work needed to disrupt these intermolecular attractions and pull apart molecules to a certain extent of area.

As the surface energy is applied to the matter to stretch its surface, the surface of the matter counteracts the surface increase through the tangential tension force. This isotropic surface stress associated with deformation is called surface tension. The terms “surface tension” and “surface energy” refer to the same dimensional quantity, as can be shown in the following analysis, although they may not necessarily be of the same value:

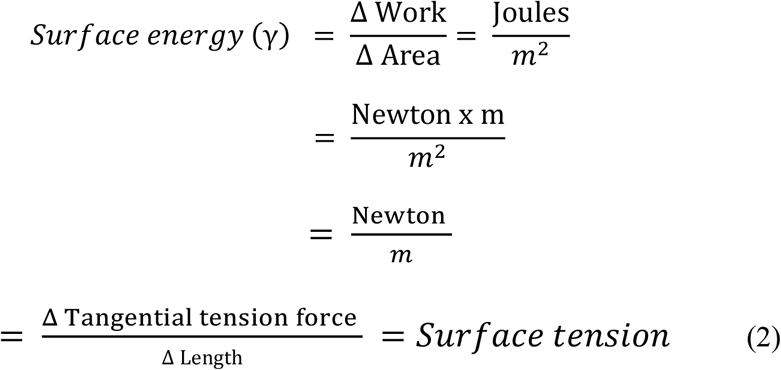

For liquid, the two values are the same because as the surface deforms, the intermolecular distance at the surface does not change because the molecules at the bulk can freely move to the deforming surface. As such, two terms are often used interchangeably for liquids. However, for a solid, the surface molecules remain constant, so the work required for surface deformation is dependent on the intermolecular distance. As a result, this work is not the same as the work required for creating a new surface (Mondal et al., 2015). As our paper deals with both solid and liquid surfaces, only the term “surface energy” will be used henceforth for simplicity.

The surface energy of water is one of the highest (72 J/m^2^ at room temperature and atmospheric pressure) because of a strong hydrogen bond between water molecules (Hauner et al., 2017). Interestingly, in a cryoEM grid, the solid particles are primarily drawn to the liquid surface, despite the high surface energy there. Such behavior is also known as the air water interface (AWI) adsorption phenomenon. To explain this phenomenon that, at a glance, seems to defy the laws of physics, we provide time-lapsed schematics of three main stages and the mathematical equation that accounts for the area of all interfaces and the respective surface energy. The equation calculates the overall potential energy for each stage, allowing for direct comparison among the stages.

In the first stage immediately after the sample is applied to a grid, particles are within the sample’s bulk volume, suspended across a grid hole away from the AWI (Fig. 2A, left). Since the surface energy (γ) is defined as the work required to build a unit area, the work (in Joules) needed to build a surface is the product of 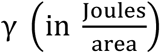 and the area created. Based on Navier-Stokes equations, which describe the motion of viscous fluid substances (Temam, 2001), the overall potential energy (Ω in Joules) for all the surfaces along the vertical axis can be calculated as the following:

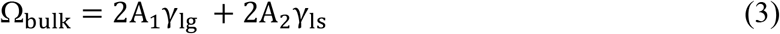

where γ_lg_ is the surface energy generated by the buffer solution (liquid) interfacing the surrounding air (gas); A_1_ is the liquid surface area, which is essentially the size of the grid hole; γ_ls_ is the surface energy at the top and bottom surfaces of the particle (solid) interfacing buffer solution (liquid); A_2_ is the top or bottom surface area of the solids, which is essentially half of the combined areas of all particles.

**Figure 2.**
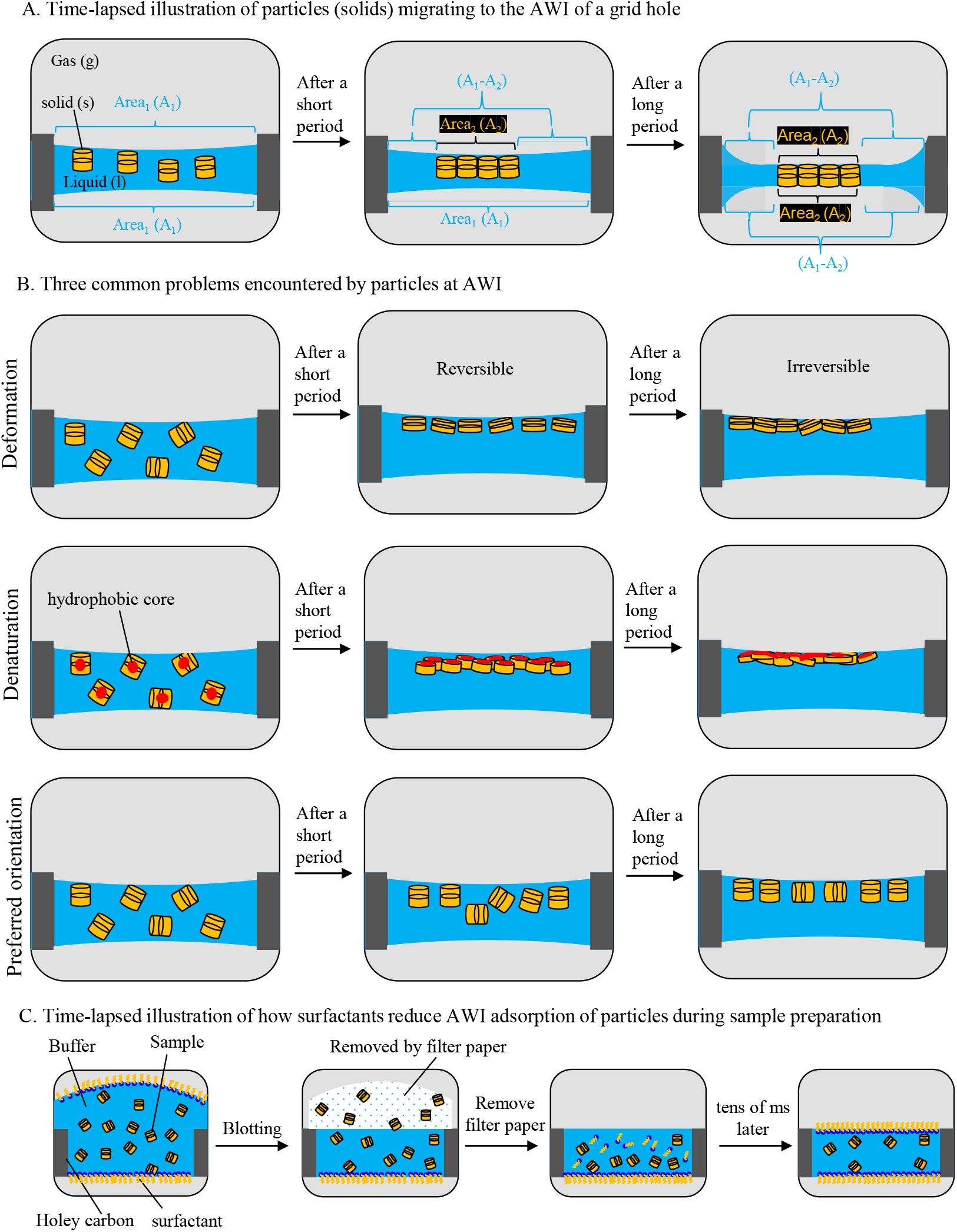
Side-view schematics of a hole of cryoEM grid, highlighting particle behavior and distribution, in relation to AWI. (A) Time-lapsed illustration of AWI adsorption of particles in the buffer. First, particles stay in the bulk volume (Left panel). A_1_ denotes the surface area of the buffer layer at the AWI, i.e., the size of the grid hole. There are two A_1_’s, one at the top and one at the bottom. After a short period of time, particles adsorb to one of the two AWIs, reaching a more favorable energy state (one-side AWI adsorption) (Center panel). A_2_ denotes the overall sum of the area of particles’ surfaces exposed to the air, upon migrating to the AWI. At the top AWI, the overall A_1_ decreases as the particles occupy the surface, unlike the unoccupied A_1_ at the bottom AWI. Eventually, particles adsorb to the remaining AWI as the buffer layer thins, reaching the most favorable energy state (two-side AWI adsorption) (Right panel). At the bottom AWI, the overall A_1_ also decreases due to the same particles occupying that surface. (B) Three common problems encountered by the sample particles at AWI, namely, particle deformation, denaturation, and preferred orientation. (C) Time-lapsed illustration of how surfactants reduce or even eliminate AWI adsorption of particles during sample preparation. First, the sample-surfactant mixture is loaded from the top side, denoted by the bulge (first panel). Surfactants migrate more quickly than the sample particle (see explanation in Section 2.3) to occupy the two AWIs, blocking sample particles from migrating to the surfaces. Upon blotting, some particles and surfactants are removed by the filter paper (second panel), exposing a second surface at the blotted side (third panel), which would also be occupied by the surfactant molecules, prevent remaining sample particles from migrating to the free surface (last panel).

In the second stage, a particle adsorbs to one of the AWIs, which we call the single adsorption state (Fig. 2A, center). The overall potential energy of the single adsorption state can be calculated as

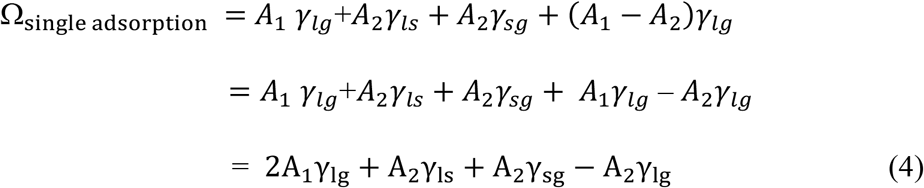

in which a new surface energy, γ_sg_, is generated by the particle’s top surface. As the particles occupy the top surface, the working surface area for γ_lg_ decreases, reducing the total work exerted by the surface energy. The change in the overall potential energy from Ω_single adsorption_ to Ω_bulk_, can be expressed as,

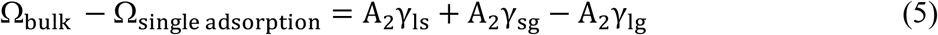

which can be further simplified using Young’s equation,

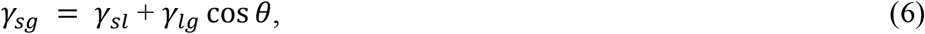

where θ is the equilibrium contact angle the liquid makes with the solid (Shuttleworth 1950, Hiemenz and Rajagopalan, 1997).

In the case of single adsorption, the contact angle between the liquid and solid is 180°, simplifying the equation to

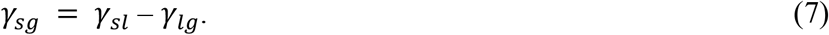

Using this relationship, the equation (5) simplifies to

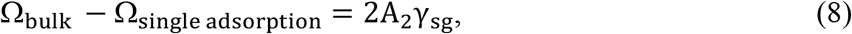

indicating a decrease in the overall potential energy as the system transitions from the bulk state to the single adsorption state.

Because the particle adsorption to the surface leads to a more energetically favorable state, the remaining particle side will likely follow the same path, leading to a double adsorption state (Fig. 2A, right). The overall potential energy of the surface of double adsorption can be expressed as

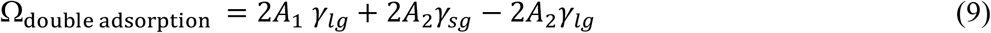

Comparing the potential energy between Ω_single adsorption_ and Ω_duble adsorption_,

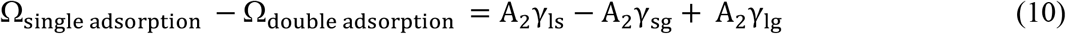

Using the relationship established in equation (7), the above difference simplifies to

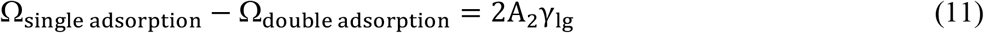

indicating a decrease in the overall potential energy as the system transitions from the single adsorption state to the double adsorption state. This explains why ice thickness of a properly blotted cryoEM grid tends to be the same as the particle thickness. Indeed, the AWI adsorption phenomenon, despite the particles’ migration to the surface, i.e., a high surface energy region, is logical as it generates the most energetically favorable state.

### 2.2 How sample aggregation is related to AWI

As commonly observed in electron microscopy, some particles aggregate at the AWI. Protein aggregation is the aftermath of protein denaturation that causes the exposed hydrophobic core of the individual proteins to come together (Koepf et al., 2017), the event prevalent at the AWI (D’Imprima et al., 2019; Wiesbauer et al., 2013). To explain the connection between AWI and protein denaturation, we adopted a coarse-grained structure-based model, which describes the force exerted on the particles at the AWI (Cieplak et al., 2014).

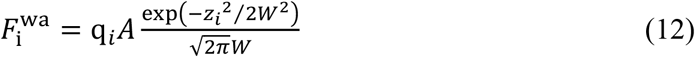

In this model, A is the amplitude of the depth of the potential in the effective contact interaction between two residues, where the higher the A of a molecule, the greater the degree of pinning of a molecule to the water surface; W is the width of the interface; *q*_*i*_ is the hydropathy index of amino acid, ranging from -4.5 for the polar arginine to 4.5 for the most hydrophobic isoleucine. The AWI is centered at z=0, where Δ z indicates protein deformation. This phenomenological model suggests that the force exerted at the AWI denatures protein, leading to the readjustment of the protein orientation to balance the hydropathy-related forces. By comparing proteins with varying degrees of hydrophilicity, others have demonstrated that the AWI adsorption occurrence is proportional to both the protein’s hydrophobic residues and the protein deformation events (Zhao et al., 2017); To equilibrate the hydropathy-related forces, the hydrophilic and hydrophobic residues of the protein pull toward the bulk and AWI, respectively, resulting in protein deformation at the AWI.

Since there are charged molecules in the sample, we also considered electrostatic forces. Toward this end, we explored the concept of Debye-Hückel length (Kirby, 2010), which measures how far the charge-carrying species’ net electrostatic effect persists in a solution. This length is defined as

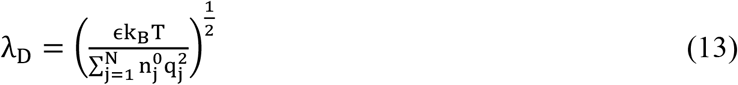

where ϵ is the relative static permittivity of solvent, k_B_ is Boltzmann’s constant, T is the temperature, n_j_ is the mean concentration of charges of the species j, and q_j_ is the charge of the species j. The Debye length in phosphate-buffered saline (PBS) at room temperature, a commonly used solution in biology and cryoEM research, is 0.7 nm (Chu et al., 2017), which is approximately the diameter of α helix secondary structure in proteins and significantly smaller than the size of folded proteins and their domains. As such, we concluded that the electrostatic force is negligible when considering molecular interactions at the AWI.

Lastly, we consider protein denaturation in relation to energy. In the energy landscape, unfolding events involve a series of small, destabilizing (uphill) steps, with small bumps (activation barriers) and dips (local minima), which are driven by the constant, reversible “sub-globally cooperative unfolding/refolding” events (Glaeser and Han, 2017; Maity et al., 2005) that occur at the molecular level. A native protein initially adsorbs to the AWI to reach a more energetically favorable state. After reaching a transitional state, i.e., the peak of the energy barrier, the protein spontaneously proceeds downhill and unfolds (Glaeser and Han, 2017). Hydrogens within the native protein can exchange with solvent as the native hydrogen bonds are transiently broken. The cytochrome c (Cytc) results show five sub-globally cooperative units, called foldons, that fold and unfold stepwise, whose amide hydrogen exchange is demonstrated by small local structural fluctuations that break one hydrogen bond at a time (Maity et al., 2005). Results from the unfolding of RNase H showed that it begins with the local fluctuations of the least stable protons on the side of the helix that is more solvent-exposed before the global unfolding. Both events place amide groups in proton exchange-competent forms in the solvent (Chamberlain et al., 1996). These transient events expose additional hydrophobic groups of the sample, causing them to adsorb to the AWI irreversibly. With no activation barrier at the surface, a spontaneous, non-reversible “catastrophic” denaturation occurs, leading to the unfolded proteins. Exposed internal regions of these unfolded proteins are typically hydrophobic, which promote sample aggregation, a hallmark of the AWI problem.

### 2.3 Surfactant application in cryoEM sample preparation alleviates AWI adsorption problem by shifting equilibrium

A surfactant is an amphipathic surface-active molecule, with its polar head in the water and non-polar tail exposed to the air. Surfactants have been widely used in cryoEM to alleviate the AWI adsorption phenomenon. Our formulation (Equation 7) predicts that all particles would migrate to the AWI as that would lower potential energy of the system (*i*.*e*., more negative Ω). It also predicts ways to solve or alleviate the AWI problem: eliminating such aqueous surfaces of the system or, if impossible, to eliminate sample particles’ access to the surface, decreasing surface energy. For the latter, surfactants can lower the surface energy (*i*.*e*., γ_lg_ above), thus rationalizing the empirical use of surfactants in alleviating the AWI problem (Liao and Zatz, 1979).

Surfactants reduce the surface energy of the aqueous solution because when surfactant molecules populate the water surface, they replace the existing hydrogen bonds between water molecules. As a result, surface energy (γ_lg_) gradually decreases until reaching the minimum at the surfactant’s critical micelle concentration (CMC), the concentration above which surfactants form micelles (Liao and Zatz, 1979). Importantly, thanks to their hydrodynamic radius, surfactants will migrate faster to the surface than the sample molecules, which typically are much larger macromolecular complexes, protecting the sample molecules from denaturation. In the extreme scenario when all sites on the AWI are occupied by the surfactant molecules, exposed surfaces available to sample molecules would be eliminated entirely, resulting in an ideal situation where aqueous surface exists for the macromolecular complexes to be imaged by cryoEM.

### 2.4 Experimental confirmation of the air-water interface adsorption problem

CryoEM sample is embedded in vitreous ice, which helps retain near-native high-resolution features of the sample. The aqueous buffer, which becomes vitreous ice through flash freezing, is a double-edged sword in that it is essential in providing a “near-native” environment for the sample, but it is its very existence that introduces AWI adsorption phenomenon, causing denaturation, deformation, and preferred orientation of the sample. To address these potential problems in cryoEM sample preparation, we observed the behavior of a widely tested non-membrane protein, GroEL. We used cryoET to reveal the 3D structure of the embedding ice in the hole of the holey carbon grids to assess the GroEL distribution throughout the grid thickness.

After generating 3D reconstructions using the IMOD package (Kremer et al., 1996), we could assess the particle distribution throughout the ice thickness, as shown in Figure 3. The slice of tomograms (about 5nm thick) was extracted from two AWIs and the middle region of the ice. Without surfactant (Fig. 3A1-3A3), most particles were observed at the AWIs, indicating that GroEL particles, like other complexes, suffer from AWI adsorption problems. More specifically, most particles were on the opposite surface (Fig. 3A1, Top slice) of the sample-applied surface (Fig. 3A3, bottom slice).

**Figure 3.**
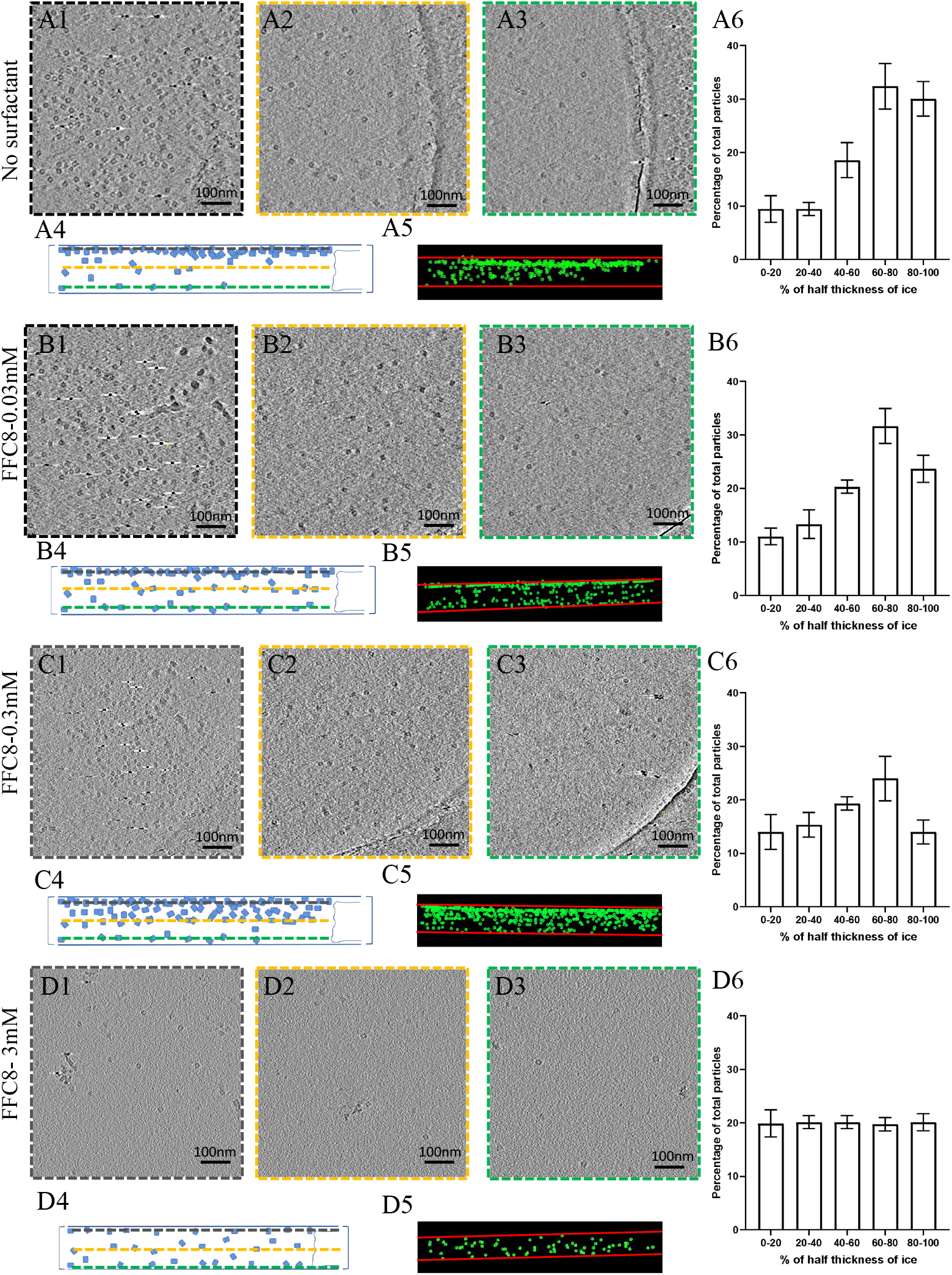
GroEL particle distribution in the vitrified sample with different concentrations of FFC8. (A-D) Each alphabetized panel (A-D) contains 6 numericized sub-sections (*e*.*g*., A1 through A6 or D1 through D6), organized in the same manner for all panels. Panels A1 through A6 pertain to no-surfactant condition, and B through D pertain to different FFC8 concentrations as indicated. The first three sub-sections (1-3) show the top, center, and bottom slice of the tomogram from cryoET, respectively, viewed from the top of the grid. Sub-section 4 shows the schematic of sub-sections (1-3), viewed from the side of the grid. The dashed border (black, orange, and green) in sub-sections 1-3 corresponds to the dashed horizontal line (black, orange, and green) in sub-section 4. Sub-section 5 shows the actual particle distribution of the tomogram, viewed from the side, with green dots representing GroEL particles and red horizontal lines representing the AWIs. Sub-section 6 shows the histogram of particle distribution across the entire ice thickness. X-axis is broken into ranges based on the magnitude of deviation (represented in %) from the center of the ice (*e*.*g*., 0 % representing no deviation from the center of the ice and 100% representing the greatest deviation from the center, i.e., either side of AWI). Y-axis represents the number of particles (represented as the percentage of the total population) residing in each ice region.

Next, we analyzed the side view from our reconstructed tomogram to better understand the particle distribution (Fig. 3A4-5). The degree of deviation away from the middle was expressed in percentage; The middle plane and the surfaces were set to 0% and 100%, respectively (Fig. 3A6). 30% of the particles were in the 80∼100% range, and 33% were in the 60∼80% range. Taken together, our results indicate that GroEL suffers from the AWI adsorption problem.

### 2.5 Different surfactants at varying concentation can alter particle distribution in ice

Upon confirming the AWI absorption problem in GroEL, we experimentally validated the effectiveness of surfactants. Our first surfactant of choice was FFC8, which can help form a thinner buffer layer for cryoEM samples (Glaeser, 2018; Hughes et al., 2018; Johnson and Chen, 2017) than the non-fluorinated counterparts. Also, FFC8 has a relatively high CMC (2.9mM) compared to many other surfactants, whose CMCs are generally less than 1mM (Inácio et al., 2011), alleviating the micelle-induced perturbation of image contrast. As a result, FFC8 tolerates a broader range of applicable concentrations. Lastly, FFC8 contains a polar tail that is both hydrophobic and oleophobic, reducing the likelihood of biomolecule denaturation (Park et al., 2007).

We prepared three samples of 2 mg/mL GroEL, each with a different concentration of FFC8 (1x CMC, 1:10 CMC, and 1:100 CMC FFC8) added right before freezing the grid, and cryoET tilt series were collected and reconstructed. The number of particles in three tomogram slices (∼5nm thick), including two slices for AWI layers and one slice for the middle layer, were compared among the samples in the above concentrations (Fig. 3B1-3B3, 3C1-3C3, 3D1-3D3). As the FFC8 concentration increased, the distribution bias toward the AWI decreased (Fig. 3B4-3B6, 3C4-3C6). At CMC, particles were evenly distributed throughout the ice thickness (Fig. 3D4-3D6). The overall particle number significantly decreased at CMC, which indicates more even particle distribution, as shown in our calculation from the previous section. Our findings are in line with the previous study that highlighted surfactants’ effectiveness in relieving the oversaturation of the field of view (Roh et al., 2017).

Aside from the fluorinated surfactant, we also tested non-ionic surfactants such as dodecyl maltoside (DDM) and NP-40, commonly used for solubilizing membrane proteins for cryoEM studies (Vinothkumar, 2015). Like FFC8 at CMC, the addition of NP-40 at CMC (0.30mM) and DDM at CMC (0.15mM) resulted in even particle distribution of GroEL (Fig. 4). These results indicate that DDM and NP40, surfactants of choice for membrane proteins, can also be effective on cytosolic proteins, such as GroEL, suggesting a broad spectrum of surfactant efficacy in improving the particle distribution. In summary, using reconstructed cryoET tomograms, we determined that the particle distribution in ice is directly related to surfactant concentration, reaching near elimination of AWI absorption phenomenon at CMC.

**Figure 4.**
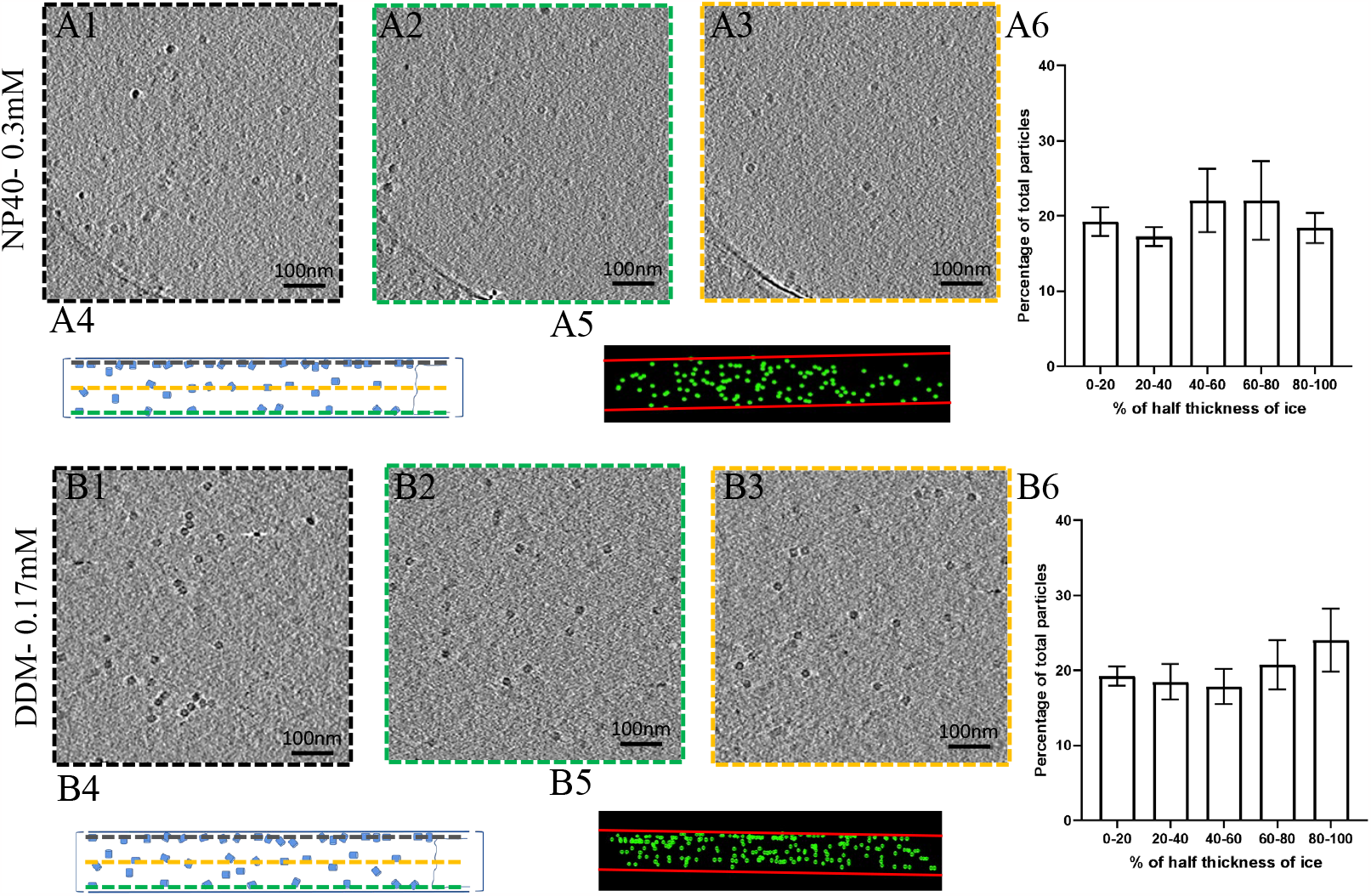
GroEL particle distributions in vitrified samples with different surfactants at their respective CMC. (A-B) Each alphabetized panel contains 6 numericized sub-sections (*e*.*g*., A1 through A6 or B1 through B6), organized in the same fashion for all panels. Panel A1 through A6 pertain to NP40 at CMC condition and B1 through B6 pertain to DDM at CMC condition as indicated. The first three sub-sections (1-3) show the top, center, and bottom slice of the tomogram from cryoET, respectively, viewed from the top of the grid. Sub-section 4 shows the schematic of sub-sections (1-3), viewed from the side of the grid. The dashed border (black, orange, and green) in sub-sections 1-3 corresponds to the dashed horizontal line (black, orange, and green) in sub-section 4. Sub-section 5 shows the actual particle distribution of the tomogram, viewed from the side, with green dots representing GroEL particles and red horizontal lines representing the AWIs. Sub-section 6 shows the histogram of particle distribution across the entire ice thickness. The X-axis is broken into ranges based on the magnitude of deviation (represented in %) from the center of the ice (*e*.*g*., 0 % representing no deviation from the center of the ice and 100% representing the greatest deviation from the center, i.e., either side of AWI). The Y-axis shows the number of particles (represented as the percentage of the total population) residing in each ice region.

### 2.6 Impact of commonly used surfactants on high-resolution cryoEM

Surfactant at high concentration is commonly used in molecular biology research to denature protein, namely, sodium dodecyl sulfate in SDS-PAGE. However, the denaturing ability of surfactants is undesirable in structural studies. To test if the surfactants harm the protein structure, we conducted a single-particle cryoEM to assess the structural integrity of GroEL in the presence of surfactants. Around 500 to 700 micrographs were collected for the conditions above (at CMC for conditions with surfactants), and 3D reconstructions from the extracted particles were compared (Fig. 5). 3D classification into 4 classes generated one “good” class average for all conditions, which was selected for the 3D refinement. The number of particles was normalized to 20,936 for all conditions. We were able to reach 3.3 Å, 3.5 Å, 3.5 Å, and 3.3 Å resolution for no-surfactant, DDM, FFC8, and NP40 conditions, respectively. Chain L of the X-ray crystal structure [PDB: 1MNF (Wang and Chen, 2003)] was fitted into the corresponding cryoEM density of all reconstructions to validate the structural integrity (Fig. 5) at the level of amino-acid side chains (bottom row of Fig. 5). Overall, these surfactants at CMC did not induce structural damage of the particles.

**Figure 5.**
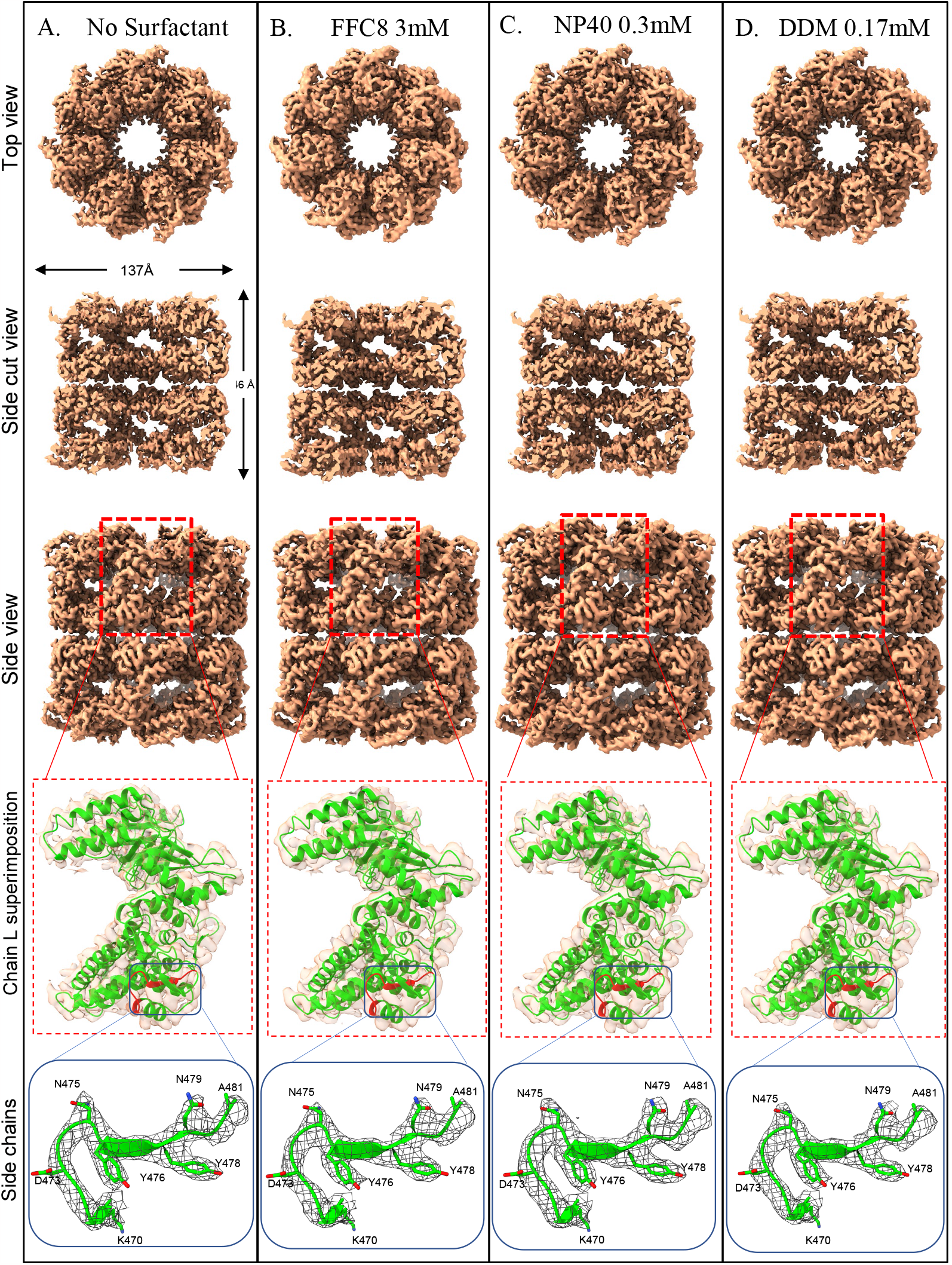
Near-atomic cryoEM reconstructions of GroEL, demonstrating no structural damage introduced by the surfactants at the indicated concentrations—the proposed solution to the AWI problem. (A-D) Various surface views of the cryoEM reconstructions of GroEL in different buffer conditions, in the absence of any surfactant (A) at 3.3 Å resolution, and the presence of 3.0 mM FFC8 (B) at 3.5 Å resolution, 0.3mM NP40 (C) at 3.3 Å resolution, and 0.17mM DDM (D) at 3.5 Å resolution. The 4^th^ row shows the semi-transparent zoom-in view of the boxed region in the 3^rd^ row, where a ribbon diagram of chain L of the X-ray crystal structure (Wang and Chen, 2003) (PDB: 1MNF, shown in green ribbon representation) was fitted. The 5^th^ row shows the zoom-in view of the boxed region in the 4^th^ row. The structure colored in red within the box (amino acid residues 470 to 481) is now shown with the amino-acid residues and the cryoEMdensities are displayed as wireframes.

Having demonstrated that surfactants can prevent AWI adsorption while maintaining the structural integrity of a stable, soluble protein, GroEL, we asked if this offers a solution to a more challenging, real-world problem, namely, ClC-1, as shown in Figure 1. ClC-1 is a membrane channel essential for maintaining Cl^-^ permeability across the plasma membrane of skeletal muscle fibers, accounting for 80% of the resting membrane conductance in humans (Stauber et al., 2012). Initially, the common phenomena that arise when the samples transition from negative-stain EM grid to cryoEM grid, namely, the emergence of aggregation and deformation of particles, were encountered by ClC-1 (Fig. 1). The application of FFC8 at CMC (3mM), before vitrification, led to mono-dispersion of ClC-1 particles (Fig. 6A), suggesting that the samples no longer migrated to the AWI, thus preventing deformation and subsequent aggregation. This step was essential in producing good quality particles and the eventual 3.6 Å overall resolution cryoEM density map of the transmembrane domain, enabling *de novo* atomic model building (Wang et al., 2019) (Fig. 6B). Taken together, surfactants application is an effective and practical solution to alleviating AWI adsorption problem in cryoEM.

**Figure 6.**
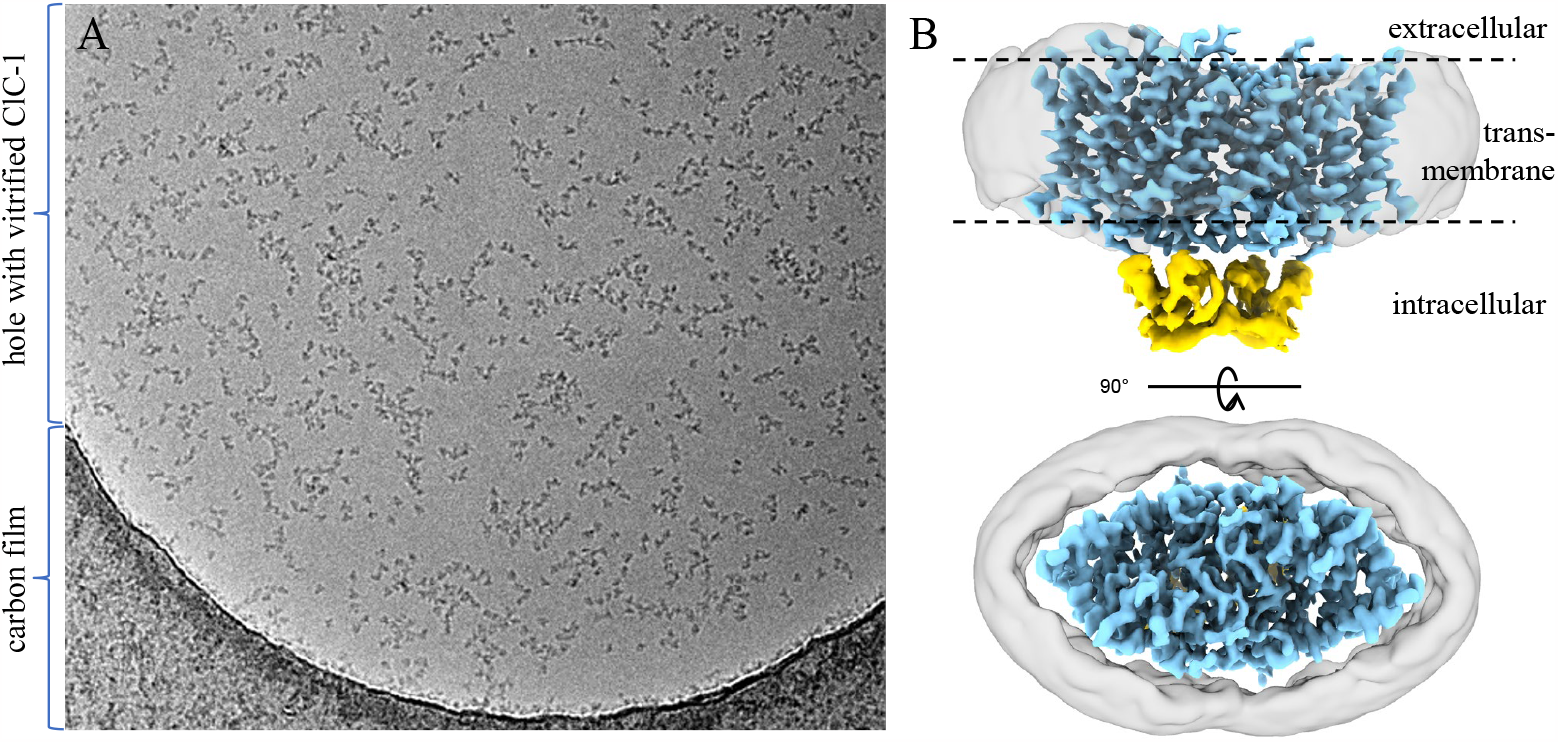
Application to real-world AWI adsorption problem. (A) A representative micrograph of single-particle cryoEM of purified human ClC-1 protein after applying FFC8 (at CMC, 3 mM). As can be compared with Figure 1, adding surfactant effectively solved the protein aggregation and deformation problem. (B) Two orthogonal shaded surface views of the resultant cryoEM map at an overall resolution of 3.6 Å, enabling *de novo* atomic modeling of the membrane domain (blue) of the complex (Wang et al., 2019). The intracellular domain is shown in yellow, and the detergent belt is in semi-transparent gray.

## 3. Discussion

As cryoEM becomes widely recognized in the structural biology community as a tool of choice for determining atomic structures of biological complexes, the cases of these complexes encountering problems of preferential orientation, aggregation, or mysterious “disappearance” on cryoEM grids—all resulting from AWI absorption phenomenon of particles— have been well-recognized. However, the reasons for such misbehavior are not well-understood, limiting systematic approaches to addressing these problems. Here, we have proposed theoretical formulations that explain the AWI adsorption phenomenon and the associated problems.

According to our formulation, the particles in the bulk solution migrate to the buffer surface to minimize the overall potential energy within the grid hole. As a result, the buffer surfaces become fully occupied by the particles. Based on this explanation, we experimentally examined the effect of surfactants application on the problems associated with the AWI adsorption phenomenon since surfactants can lower the surface energy (γ_lg_) by occupying the buffer surface to decrease the overall potential energy. To this end, we conducted cryoET to construct tomograms of the entire grid thickness to visualize the particles distribution in the hole of cryoEM grids. Indeed, our cryoET showed that all tested surfactants at CMC reduced the AWI adsorption of particles; FFC8 at CMC resulted in the most even distribution. Without surfactant, most particles were on the opposite surface to where the sample was applied. We conjecture that the particles began migrating to the opposite surface when the sample-applied surface was blotted and that most particles had already finished migrating before the grid plunged, given that the particles can adsorb to the AWI in a matter of 0.1 s (Naydenova and Russo, 2017; Noble et al., 2018b; Taylor and Glaeser, 2008). Subsequent single-particle cryoEM confirmed that adding surfactant at CMC does not adversely affect the GroEL structure, as shown by near-atomic resolution reconstructions that are on par with that of the no-surfactant particles.

A question that arises naturally is what happens to the particles in the bulk volume, such as in the Eppendorf tube during long-term storage. Our theory suggests that particles would migrate to the surface until the surface is fully occupied by these particles. At this point, given the much smaller number of sites available on the surface area as compared to the bulk volume, the remaining particles would stay in the bulk volume, essentially eliminating the AWI problem for these particles. That is why the AWI adsorption problem has escaped observation until the advent of cryoEM, in which the particles are exposed to the greater surface and smaller bulk volume, a situation created by sample blotting. Our study rationalizes using surfactants in cryoEM sample preparation and experimentally demonstrates its effectiveness in lowering the surface energy (γ_lg_) and alleviating the AWI adsorption problem. However, potential concerns associated with the use of surfactants, such as micelle formation that can increase background noise and particle denaturation, remain. Thus, other alternatives to surfactants are also worth exploring. For one, introducing a layer of 2D particle arrays on the AWI can be a barrier to protect other particles from AWI absorption (Glaeser and Han, 2017). Another alternative is the new class of surfactants called amphipols (Liao et al., 2013; Tribet et al., 1996), which enable handling membrane proteins in a detergent-free solution. Recently, a support film using 2D crystals of hydrophobin HFBI was developed. The hydrophilic side of HFBI adsorbs sample particles via electrostatic interactions to protect them from the AWI, enabling thin ice formation for enhanced data collection (Fan et al., 2021). Other advanced engineering approaches aim to improve the cryoEM result quality without surfactants. For example, novel equipment is developed to reduce the time before the sample plunges into liquid ethane (Noble et al., 2018a), and novel support film (Han et al., 2020), like monolayer graphene-supporting film and modified graphene/graphene oxides, is developed to improve cryoEM sample quality. More recently, *in situ* structures of ribosomes and expressomes were solved inside microbial cells via cryoET (Hoffmann et al., 2022; O’Reilly et al., 2020; Tegunov et al., 2021), avoiding the AWI problem altogether. However, at the time of this writing, such successes are still limited to large complexes exemplified by ribosomes or ribosome-containing super-complexes; as such, high-resolution structures will continue to depend on practical solutions to resist or level the potential energy at AWI, as reported here both through theoretical consideration and experimental demonstration.

## 4. Materials and methods

### 4.1 Sample preparation

Chaperonin 60 lyophilized powder (GroEL, 1 mg) from *Escherichia coli* was purchased from Sigma-Aldrich (Cat # C7688, Sigma-Aldrich, St. Louis, MO, USA). GroEL powder was solubilized in 50mM Tris-HCl (pH7.5), 10mM KCl, an 10mM MgCl_2_. Subsequently, GroEL concentration was determined by NanoDrop Microvolume UV-Vis Spectrophotometers based on Protein A280. For short-term storage, 2mg/mL GroEL was stored at 4 °C and used within 2 weeks. For long-term storage, stock solutions were stored at -80 °C and thawed before use.

### 4.2 Grids preparation for cryoEM

GroEL solution was prepared as 2mg/mL and mixed with 1:70 diluted 5-nm diameter fiducial gold beads in an 8:1 volume ratio. FEI Vitrobot Mark IV was used to make vitrified samples. The sample was applied to the carbon side of 200 mesh Cu Quantifoil 100 holey carbon films (R 2/1), which were beforehand glow discharged by Gatan Plasma System SOLARUS. Tested surfactants (concentrations shown in Table 1) were directly mixed with the sample. The mixture was applied to the grid. The grids were blotted with filter paper to remove the extra sample and then plunged into liquid ethane. Grids were stored in liquid nitrogen.

**Table 1.** List of tested surfactants and their concentrations.

| 669 | Surfactant name | Surfactant concentration |  |  |
| --- | --- | --- | --- | --- |
| 670 | No surfactant | Non-applicable |  |  |
| 671 | fluorinated fos-choline 8 | 2.9mM | 0.29mM | 0.029mM |
| 672 | DDM | 0.15mM |  |  |
| 673 | NP40 | 0.3mM |  |  |

### 4.3 Single-particle cryoEM data collection

The grids were loaded into a Titan Krios electron microscope (Thermo Fisher Scientific) equipped with a Gatan imaging filter (GIF), and cryoEM images were recorded on a post-GIF Gatan K3 Summit direct electron detection camera operated in super-resolution electron-counting mode. The magnification was 81,000 ×, with a pixel size of 1.1038 Å/pixel at the specimen level. Data collection was facilitated by SerialEM (Mastronarde, 2005). The dosage rate was set to 30 electrons/Å2 on the sample level, and the exposure time for each frame was 0.2 seconds. The targeted under-focus value was 1.8 μm – 2.2 μm. In total, 500-700 micrographs were collected.

### 4.4 Single-particle cryoEM data processing

We followed the computational steps under the framework of WARP (Tegunov and Cramer, 2019) for the preprocessing, which included motion correction for the frame alignment and estimation of local defocus and resolution. Particle-picking was performed based on a machine-learning algorithm (BoxNet) in WARP, generating, on average, 173,000 extracted particles with a box size of 400 pixels (pixel size: 1.1038 Å) for all conditions. The extracted particles were classified with the “2D Classification” tool on the Relion GUI with the following parameters: Number of classes: 32, Regularisation parameter T: 2, Number of iterations: 25, Mask diameter: 300 Å. After 2D classification, “good” class averages with clear top/side views of GroEL, as well as those that fit the general dimensions of GroEL were manually selected for 3D classification, using “Subset selection” tool on the Relion GUI. The 3D classification was conducted with the following parameters: Symmetry: D7, Number of classes: 4, Regularisation parameter T: 4, Number of iterations: 25, Mask diameter: 230 Å. After 3D classification, “good” class averages were manually selected for 3D refinement using the “Subset selection” tool on the Relion GUI. Particle images belonging to the selected “good” class averages were normalized to 20,936 particles. The normalized particles were subjected to a D7 symmetry refinement with a mask diameter of 200 Å, using “3D auto-refine” on the Relion GUI. Subsequent fitting of the crystal structure and images were created in Chimera X (Goddard et al., 2018).

### 4.5 CryoET data collection

Single particle micrographs were collected on the FEI Tecnai TF20 at 200 kV equipped with a TIETZ F415MP 16-megapixel CCD camera. TEM Imaging & Analysis (TIA) was used to acquire data. All tilt series were collected from -50° to +50° with 2° increments using FEI TEM Batch Tomography software with -6 μm nominal defocus. The cumulative dose count was 50∼60 e-/Å2 per tilt series. The pixel size was 4.4 Å/pixel at the specimen level with the 50,000× nominal magnification for imaging.

### 4.6 CryoET data processing

The tilt series were reconstructed by the Etomo component of the IMOD software package (Kremer et al., 1996) to 3D tomograms. “Build Tomogram” was chosen to start the reconstruction process. Tilt series images were pre-processed by removing the outlier pixel values in the data files. Coarse alignment was done using the fiducial seeding model, followed by a fine alignment. 10 to 15 gold beads were picked for each tilt series, and a mathematical model for specimen movements was used to predict the gold beads’ positions. The mean residual error was reduced during fine alignment by fixing big residuals. Positioning tomogram thickness was set to 1000 nm to include the top and bottom AWI. The tilt axis and Z shift were also computed and adjusted to create the final alignment. The final tomograms were built using SIRT with 6 iterations. Finally, the whole tomogram was rotated around the x-axis to make the air-water interface roughly parallel to the field of view.

GroEL particles were selected and saved as a mod file. Three points on the same plane were used to identify the AWI and the middle plane. The coordinates of all particles were recorded and used to calculate the distance to each plane. Particle distribution was visualized as a 3D model by IMOD.

## Acknowledgments

We thank Ke Ding for the discussion during the planning phase of this study, Ivo Atanasov for cryoEM technical support, and Caiyan Wang for initial imaging efforts. This research was supported in part by a grant from the US National Institutes of Health (R01GM071940). We acknowledge the use of resources in the Electron Imaging Center for Nanomachines supported by UCLA and grants from NIH (S10RR23057, S10OD018111, and U24GM116792) and NSF (DBI-1338135 and DMR-1548924).

## Author Contributions

ZHZ initialized and supervised the project; XZ performed cryoET, and JSK and XZ processed the data; JSK, XZ, YL, and KW performed cryoEM imaging; JSK performed single-particle reconstruction; ZHZ, JSK, and XZ wrote the paper; all authors edited and approved the paper.

## Competing Interests

The authors declare no competing interests.

## Notes

### Competing Interest Statement

The authors have declared no competing interest.

### Summary of Updates

We revised several sentences and other minor details.

## References

Bai, X.C., Fernandez, I.S., McMullan, G., Scheres, S.H., 2013. Ribosome structures to near-atomic resolution from thirty thousand cryo-EM particles. Elife 2, e00461.

Chamberlain, A.K., Handel, T.M., Marqusee, S., 1996. Detection of rare partially folded molecules in equilibrium with the native conformation of RNaseH. Nat Struct Biol 3, 782–787.

Chen, J., Noble, A.J., Kang, J.Y., Darst, S.A., 2019. Eliminating effects of particle adsorption to the air/water interface in single-particle cryo-electron microscopy: Bacterial RNA polymerase and CHAPSO. Journal of Structural Biology: X 1, 100005.

Chen, S., Li, J., Vinothkumar, K.R., Henderson, R., 2022. Interaction of human erythrocyte catalase with air-water interface in cryoEM. Microscopy 71, i51–i59.

Chu, C.H., Sarangadharan, I., Regmi, A., Chen, Y.W., Hsu, C.P., Chang, W.H., Lee, G.Y., Chyi, J.I., Chen, C.C., Shiesh, S.C., Lee, G.B., Wang, Y.L., 2017. Beyond the Debye length in high ionic strength solution: direct protein detection with field-effect transistors (FETs) in human serum. Sci Rep 7, 5256.

Cieplak, M., Allan, D.B., Leheny, R.L., Reich, D.H., 2014. Proteins at Air–Water Interfaces: A Coarse-Grained Model. Langmuir 30, 12888–12896.

D’Imprima, E., Floris, D., Joppe, M., Sánchez, R., Grininger, M., Kühlbrandt, W., 2019. Protein denaturation at the air-water interface and how to prevent it. eLife 8, e42747.

Dubochet, J., Adrian, M., Chang, J.J., Homo, J.C., Lepault, J., McDowall, A.W., Schultz, P., 1988. Cryo-electron microscopy of vitrified specimens. Q. Rev. Biophys. 21, 129–228.

Fan, H., Wang, B., Zhang, Y., Zhu, Y., Song, B., Xu, H., Zhai, Y., Qiao, M., Sun, F., 2021. A cryo-electron microscopy support film formed by 2D crystals of hydrophobin HFBI. Nature Communications 12, 7257.

Glaeser, R.M., 2018. Proteins, Interfaces, and Cryo-Em Grids. Curr Opin Colloid Interface Sci 34, 1–8.

Glaeser, R.M., Han, B.-G., 2017. Opinion: hazards faced by macromolecules when confined to thin aqueous films. Biophysics Reports 3, 1–7.

Goddard, T.D., Huang, C.C., Meng, E.C., Pettersen, E.F., Couch, G.S., Morris, J.H., Ferrin, T.E., 2018. UCSF ChimeraX: Meeting modern challenges in visualization and analysis. Protein Sci 27, 14–25.

Graham, D.E., Phillips, M.C., 1979. Proteins at liquid interfaces: I. Kinetics of adsorption and surface denaturation. Journal of Colloid and Interface Science 70, 403–414.

Han, Y., Fan, X., Wang, H., Zhao, F., Tully, C.G., Kong, J., Yao, N., Yan, N., 2020. High-yield monolayer graphene grids for near-atomic resolution cryoelectron microscopy. Proc Natl Acad Sci U S A 117, 1009–1014.

Hauner, I.M., Deblais, A., Beattie, J.K., Kellay, H., Bonn, D., 2017. The Dynamic Surface Tension of Water. The Journal of Physical Chemistry Letters 8, 1599–1603.

Hiemenz, P., Rajagopalan, R., 1997. Principles of Colloid and Surface Chemistry, Revised and Expanded 3^rd^ Edition. CRC press.

Hoffmann, P.C., Kreysing, J.P., Khusainov, I., Tuijtel, M.W., Welsch, S., Beck, M., 2022. Structures of the eukaryotic ribosome and its translational states in situ. Nature Communications 13, 7435.

Hughes, T.E.T., Lodowski, D.T., Huynh, K.W., Yazici, A., Del Rosario, J., Kapoor, A., Basak, S., Samanta, A., Han, X., Chakrapani, S., Zhou, Z.H., Filizola, M., Rohacs, T., Han, S., Moiseenkova-Bell, V.Y., 2018. Structural basis of TRPV5 channel inhibition by econazole revealed by cryo-EM. Nat Struct Mol Biol 25, 53–60.

Inácio, Â.S., Mesquita, K.A., Baptista, M., Ramalho-Santos, J., Vaz, W.L.C., Vieira, O.V., 2011. In vitro surfactant structure-toxicity relationships: implications for surfactant use in sexually transmitted infection prophylaxis and contraception. PLoS One 6, e19850–e19850.

Jain, T., Sheehan, P., Crum, J., Carragher, B., Potter, C.S., 2012. Spotiton: a prototype for an integrated inkjet dispense and vitrification system for cryo-TEM. J Struct Biol 179, 68–75.

Johnson, Z.L., Chen, J., 2017. Structural Basis of Substrate Recognition by the Multidrug Resistance Protein MRP1. Cell 168, 1075–1085 e1079.

Kirby, B.J., 2010. Micro- and Nanoscale Fluid Mechanics: Transport in Microfluidic Devices Cambridge University Press.

Koepf, E., Schroeder, R., Brezesinski, G., Friess, W., 2017. The film tells the story: physical-chemical characteristics of IgG at the liquid-air interface. European Journal of Pharmaceutics and Biopharmaceutics 119, 396–407.

Kremer, J.R., Mastronarde, D.N., McIntosh, J.R., 1996. Computer visualization of three-dimensional image data using IMOD. J Struct Biol 116, 71–76.

Liao, M., Cao, E., Julius, D., Cheng, Y., 2013. Structure of the TRPV1 ion channel determined by electron cryo-microscopy. Nature 504, 107–112.

Liao, W.-C., Zatz, J.L., 1979. Surfactant solutions as test liquids for measurement of critical surface tension. Journal of Pharmaceutical Sciences 68, 486–488.

Lyumkis, D., 2019. Challenges and opportunities in cryo-EM single-particle analysis. J Biol Chem 294, 5181–5197.

MacRitchie, F., 1985. Desorption of proteins from the air/water interface. Journal of Colloid and Interface Science 105, 119–123.

Maity, H., Maity, M., Krishna, M.M.G., Mayne, L., Englander, S.W., 2005. Protein folding: The stepwise assembly of foldon units. Proceedings of the National Academy of Sciences 102, 4741–4746.

Marcus, P., Protopopoff, É., Maurice, V., 2019. 5 - Surface Chemistry and Passivation of Metals and Alloys, p. 91–120, in: C. Blanc and I. Aubert, Eds.), Mechanics - Microstructure - Corrosion Coupling, Elsevier.

Mastronarde, D.N., 2005. Automated electron microscope tomography using robust prediction of specimen movements. J Struct Biol 152, 36–51.

Mondal, S., Phukan, M., Ghatak, A., 2015. Estimation of solid-liquid interfacial tension using curved surface of a soft solid. Proc Natl Acad Sci USA 112, 12563–12568.

Narsimhan, G., Uraizee, F., 1992. Kinetics of adsorption of globular proteins at an air-water interface. Biotechnology Progress 8, 187–196.

Naydenova, K., Russo, C.J., 2017. Measuring the effects of particle orientation to improve the efficiency of electron cryomicroscopy. Nature Communications 8, 629.

Noble, A.J., Wei, H., Dandey, V.P., Zhang, Z., Tan, Y.Z., Potter, C.S., Carragher, B., 2018a. Reducing effects of particle adsorption to the air-water interface in cryo-EM. Nat Methods 15, 793–795.

Noble, A.J., Dandey, V.P., Wei, H., Brasch, J., Chase, J., Acharya, P., Tan, Y.Z., Zhang, Z., Kim, L.Y., Scapin, G., Rapp, M., Eng, E.T., Rice, W.J., Cheng, A., Negro, C.J., Shapiro, L., Kwong, P.D., Jeruzalmi, D., des Georges, A., Potter, C.S., Carragher, B., 2018b. Routine single particle CryoEM sample and grid characterization by tomography. Elife 7.

O’Reilly, F.J., Xue, L., Graziadei, A., Sinn, L., Lenz, S., Tegunov, D., Blotz, C., Singh, N., Hagen, W.J.H., Cramer, P., Stulke, J., Mahamid, J., Rappsilber, J., 2020. In-cell architecture of an actively transcribing-translating expressome. Science 369, 554–557.

Pantelic, R.S., Suk, J.W., Magnuson, C.W., Meyer, J.C., Wachsmuth, P., Kaiser, U., Ruoff, R.S., Stahlberg, H., 2011. Graphene: Substrate preparation and introduction. J Struct Biol 174, 234–238.

Park, K.H., Berrier, C., Lebaupain, F., Pucci, B., Popot, J.L., Ghazi, A., Zito, F., 2007. Fluorinated and hemifluorinated surfactants as alternatives to detergents for membrane protein cell-free synthesis. Biochem J 403, 183–187.

Roh, S.-H., Hryc, C.F., Jeong, H.-H., Fei, X., Jakana, J., Lorimer, G.H., Chiu, W., 2017. Subunit conformational variation within individual GroEL oligomers resolved by Cryo-EM. Proceedings of the National Academy of Sciences 114, 8259–8264.

Russo, C.J., Passmore, L.A., 2014. Controlling protein adsorption on graphene for cryo-EM using low-energy hydrogen plasmas. Nat Methods 11, 649–652.

Russo, C.J., Passmore, L.A., 2016. Progress towards an optimal specimen support for electron cryomicroscopy. Curr Opin Struct Biol 37, 81–89.

Shuttleworth, R., 1950. The Surface Tension of Solids. Proceedings of the Physical Society 63, 444–457.

Stauber, T., Weinert, S., Jentsch, T.J., 2012. Cell biology and physiology of CLC chloride channels and transporters. Compr Physiol 2, 1701–1744.

Taylor, K.A., Glaeser, R.M., 2008. Retrospective on the early development of cryoelectron microscopy of macromolecules and a prospective on opportunities for the future. Journal of Structural Biology 163, 214–223.

Tegunov, D., Cramer, P., 2019. Real-time cryo-electron microscopy data preprocessing with Warp. Nat Methods 16, 1146–1152.

Tegunov, D., Xue, L., Dienemann, C., Cramer, P., Mahamid, J., 2021. Multi-particle cryo-EM refinement with M visualizes ribosome-antibiotic complex at 3.5 A in cells. Nat Methods 18, 186–193.

Temam, R., 2001. Navier-Stokes Equations: Theory and Numerical Analysis AMS Chelsea Pub.

Tribet, C., Audebert, R., Popot, J.-L., 1996. Amphipols: Polymers that keep membrane proteins soluble in aqueous solutions. Proceedings of the National Academy of Sciences 93, 15047.

Vinothkumar, K.R., 2015. Membrane protein structures without crystals, by single particle electron cryomicroscopy. Current opinion in structural biology 33, 103–114.

Wang, J., Chen, L., 2003. Domain Motions in GroEL upon Binding of an Oligopeptide. Journal of Molecular Biology 334, 489–499.

Wang, K., Preisler, S.S., Zhang, L., Cui, Y., Missel, J.W., Gronberg, C., Gotfryd, K., Lindahl, E., Andersson, M., Calloe, K., Egea, P.F., Klaerke, D.A., Pusch, M., Pedersen, P.A., Zhou, Z.H., Gourdon, P., 2019. Structure of the human ClC-1 chloride channel. PLoS Biol 17, e3000218.

Wiesbauer, J., Prassl, R., Nidetzky, B., 2013. Renewal of the Air–Water Interface as a Critical System Parameter of Protein Stability: Aggregation of the Human Growth Hormone and Its Prevention by Surface-Active Compounds. Langmuir 29, 15240–15250.

Williams, R.C., Glaeser, R.M., 1972. Ultrathin carbon support films for electron microscopy. Science 175, 1000–1001.

Zhao, Y., Chwastyk, M., Cieplak, M., 2017. Topological transformations in proteins: effects of heating and proximity of an interface. Scientific Reports 7, 39851.

